# Fishing the line depends on reserve benefits, individual losing at boundary and movement preference

**DOI:** 10.1101/2020.09.15.299032

**Authors:** Renfei Chen, Marissa L. Baskett, Alan Hastings

## Abstract

Whether fishing around the marine reserve edge can enhance harvested yields is an important issue in fisheries management. To solve the conundrum is difficult because of the lack of a matched boundary condition. Here, we derive a new boundary condition by considering individual losing at habitat boundaries. With the suitable boundary condition, our results suggest that individuals with high growth rate inside but low growth rate outside the reserve and high movement preference to a large marine reserve boundary can enhance yields benefits from fishing around the marine reserve edge. The findings provide theoretical cautions for fishing near some new reserves in which population growth rate might be low. Moreover, our boundary condition is general enough for the universal phenomenon of losing individual at habitat boundaries such as being applied into classic theories in refuge design to explain some previous counter-intuitive phenomena more reasonably.

## Introduction

Human activities such as over-exploiting marine resources in fisheries have entailed declines in marine populations and altered marine communities, which may further affect marine ecosystem-level properties and the sustainable delivery of marine ecosystem services [1]. An effective approach to achieve marine conservation goal is restricting human activities, and increasing marine reserves and/or marine protected areas are established since the 1980s [1]. Indeed, researches have shown that marine reserves could enhance ecological resilience[2]. On the other hand, fishers want to achieve high harvested yields. Thus, a central issue in ecology is that whether the implementation of marine reserves can meet multiple goals by improving harvested yields in fisheries management as well as maintaining population persistence in species conservation. Theoretical frameworks indicate that marine reserves can produce equivalent harvested yields in comparison with traditional fisheries management without a marine reserve [3], and the reserve benefits in improving the harvested yields of target species still exist even if considering the persistence of the other easily endangered species [4]. However, the establishment of marine reserves also gives rise to subsequent changes in a fisher’s behavior whose consequences on fisheries management still not clear. For example, if a certain marine area is closed, fishermen tend to use the tactic of “fishing the line”: concentrating fishing effort near the boundary of a marine reserve [5, 6, 7].

The universal harvesting strategy of fishing the line is based on the principle of spillover effect which suggests that the net export of adult fish from the marine reserve can benefit the adjacent fishing areas in fisheries yields [6, 8, 9]. Given anticipated higher densities at reserve boundaries, fishing the line might increase an individual fisher’s catch per unit effort. Such a response might lead to an increased harvested yield per unit coastline length. Moreover, the hypothesis of fishing the line has been supported by empirical data [10]. With long-term surveying data before and after the implementation of marine protected areas at the California Channel Islands, Cabral et al. (2016) found that commercial dive boats prefer to stay near the boundaries of marine protected areas after it was established. Nevertheless, the advantages of fishing the line coexist with some disadvantages to the unprotected harvested area. Compress fishing effort into small areas near a marine reserve results in large individual mortality there, which may erode the reserve benefits such as spillover of adults and larvae dispersal to the harvested areas and thus the total harvested yields in the harvested area could probably decrease. Moreover, the advantages of fishing the line in probably increasing fish catch per unit effort does not mean increasing the total potential harvested yields at the boundary. Thus, whether fishing the line could produce higher total harvested yields than fishing in the harvested area is still not clear. To address this issue, a boundary condition with discontinuities in both flux and density must be derived theoretically. This derives from the fact that the harvesting tactic of fishing the line could cause large amount of fish individuals losing and an abrupt density change at habitat boundaries.

An integrated theoretical framework benefits to understand how boundary conditions regulate population dynamics and persistence. Recently, a theoretical model, including both continuous and discontinuous density, makes an important step on that direction [11, 12, 13]. Its generality makes the integrated boundary condition extremely applicable to many areas such as refuge design and reserve networks [11, 14, 15]. However, the integrated model is based on the assumption of the principle of flux continuity, which implies that all individuals who leave one patch must enter another without any losses or additions of individuals at the boundary [11, 15]. This assumption prevents applying the integrated boundary condition into the harvesting tactic of fishing the line in which fish individuals are lost at the reserve boundary.

In this paper, we focus on the issue that studying the advantages of fishing the line in harvested yields under appropriate boundary conditions. First, we start by mathematically deriving a boundary condition with considering fish individual losing at habitat boundaries, which can be applied into the harvesting tactic of fishing the line. In addition to being suitable for the scenario that individual losing at habitat boundaries, our boundary conditions will be a more integrated boundary condition. With generality, our boundary conditions can turn into previous boundary conditions under special cases. We can achieve boundary conditions derived by Maciel and Lutscher (2013) with discontinuous density and continuous flux with two additional assumptions: 1) no individuals are lost at habitat boundaries, and the movement probabilities in each patch are the same (or the same step sizes in each patch); 2) individuals move from the boundary into certain patch with the step size that equals the average step size (or the average movement probability) in both patches multiplied by the same movement probability (or the same step size) in both patches. Second, we develop a theoretical framework which shows that the fishing effort per unit length at the reserve boundary is much strong and that the population persistence in the harvested area may not depend on the adult spillover from the marine reserve. Our new boundary conditions are then applied into the theoretical framework to make a comparison in the harvested yields between fishing in the harvested area and fishing the line so that to investigate the advantages and limitations of fishing the line. To make that further, we study how the relative yields (i.e. the quotient of the yields from harvested area and the yields from fishing the line) is regulated by marine reserve benefits (high growth rate in a large marine reserve), individual movement behavior as well as individuals losing at the reserve boundary. Third, to discuss the applications of our new boundary conditions, we apply them into previous issues in refuge design to observe whether the counter-intuitive results suggested in previous research still exist. By reanalyzing the existed model in refuge design, we will show the more reasonable explanations that our boundary conditions might give to ecological issues in conservation.

## Methods

We begin with a one-dimensional model described by the Fisher-KPP equation [16, 17]. With the addition of fishing effort, Herrera *et al*. used this equation to study the non-cooperative Cournot–Nash equilibrium for fishing harvest in high seas among different states [18]. Here, we only study one group of fishers in the harvested area, while there is no anthropogenic fishing activities in the marine reserve. Between the harvested area and the marine reserve, we consider a narrow band with elevated and intensive fishing efforts. The narrow band will become a boundary when its size approaches zero so that fishing in the band turns into the so-called “fishing the line”. We normalise the coastline length of a marine system to be 1. Among which, the coastline length of the marine reserve is *L*, and the coastline length of the narrow band is *ε*. Thus, the interior boundaries occurring at the position of *L* and *L* + *ε* (see Fig. 1). We define *D*_*i*_ as diffusion rate in patch *i* (*i* = 1 for harvested area, *i* = 2 for marine reserve, *i* = 3 for narrow band). At location *x* and time *t*, the dynamics of population density *u*_*i*_ in patch *i* is

**Figure 1:**
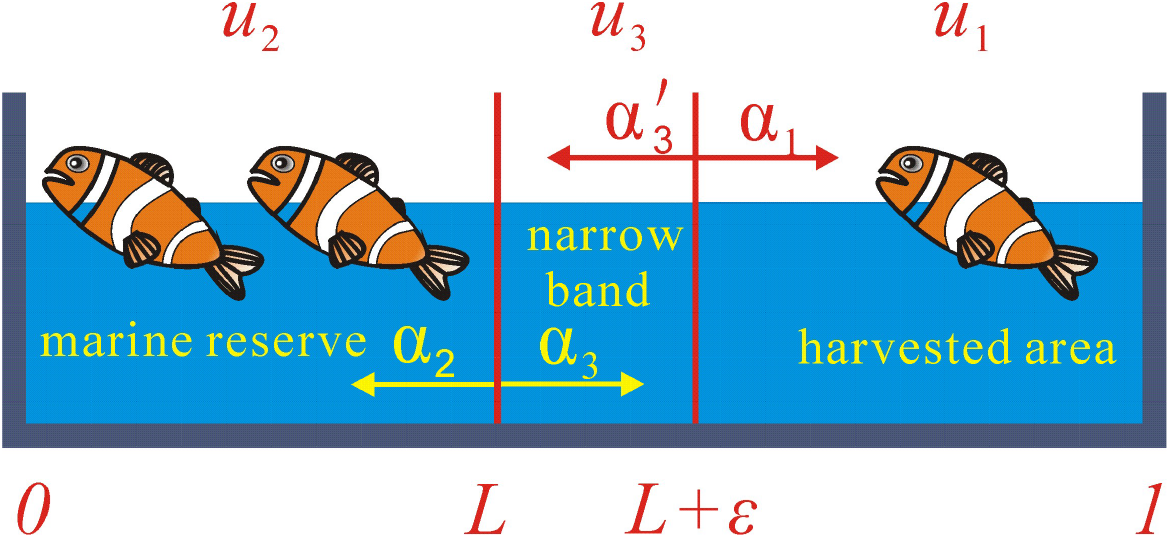
Schematic diagram of a marine system for fisheries with a total coastline length 1. The system consists of the marine reserve, the narrow band and the harvested area. The coastline length of the marine reserve and the narrow band is *L* and *ε*, respectively. *u*_1_, *u*_2_ and *u*_3_ are the density in the harvested area, in the marine reserve and in the narrow band, respectively. *α*_1_, *α*_2_ and *α*_3_ 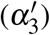 are the individual moving probability at the boundary toward the harvested area, the marine reserve and the narrow band, respectively.

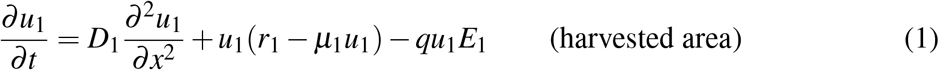

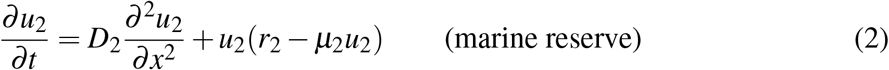

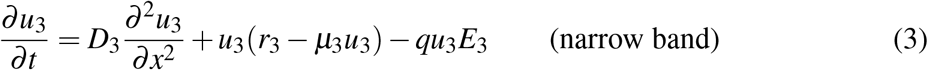

where *q* is the catch-ability rate which was assumed the same both in the harvested area and in the narrow band. *r*_*i*_ is the intrinsic growth rates, and *µ*_*i*_ measures the non-linear effects. *E*_1_ is fishing effort per unit length in the harvested area. *E*_3_ is fishing effort per unit length in the narrow band, which represents the intensive fishing activities in the narrow band and is much larger than *E*_1_ (see more definitions of symbols in table 1). We assume that the product of fishing effort *E*_3_ and the coastline length in the narrow band *ε* is a constant *β* (i.e. *εE*_3_ = *β*). According to this assumption, the maximum value of fishing effort per unit length in the narrow band corresponds to the minimum value of coastline length in the narrow band, and *E*_3_ will be infinite if *ε* approaches zero. Therefore, this assumption can fit the phenomenon of fishing the line.

**Table 1:** Definitions of the variables and parameters in the paper.

| Symbol | Definition |
| --- | --- |
| $L$ | Coastline length of the marine reserve. |
| $\varepsilon$ | Coastline length of the narrow band. |
| $\beta$ | Total fishing effort in the narrow band. |
| $u_1$ | Density in the harvested area. |
| $u_2$ | Density in the marine reserve. |
| $u_3$ | Density in the narrow band. |
| $r_1$ | Intrinsic growth rate in the harvested area. |
| $r_2$ | Intrinsic growth rate in the marine reserve. |
| $r_3$ | Intrinsic growth rate in the narrow band. |
| $D_1$ | Diffusion rate in the harvested area. |
| $D_2$ | Diffusion rate in the marine reserve. |
| $D_3$ | Diffusion rate in the narrow band. |
| $k$ | Density discontinuity at the boundary between the marine reserve and the harvested area. |
| $k_1$ | Density discontinuity at the boundary between the narrow band and the harvested area. |
| $k_2$ | Density discontinuity at the boundary between the narrow band and the marine reserve. |
| $m$ | Flux discontinuity at the boundary between the marine reserve and the harvested area. |
| $m_1$ | Flux discontinuity at the boundary between the narrow band and the harvested area. |
| $m_2$ | Flux discontinuity at the boundary between the narrow band and the marine reserve. |
| $E_1$ | Fishing effort per unit length in the harvested area. |
| $E_3$ | Fishing effort per unit length in the narrow band. |
| $q$ | Catch-ability rate. |
| $Y_1$ | Harvested yields in the harvested area. |
| $Y_3$ | Harvested yields in the narrow band<br>(i.e. yields from fishing the line when $\varepsilon$ approaches zero). |

By generalizing previous work on boundary conditions, we model individual movement with habitat preference at the boundary between two different patches and with random walk within each patch. We define *p*_*i*_ as movement probability in patch *i* with random walk (Recall that *i* = 1 for harvested area, *i* = 2 for marine reserve, *i* = 3 for narrow band). With habitat preference, individuals at position *L* move to the narrow band with probability *α*_3_ and to the marine reserve with probability *α*_2_. Individuals at position *L* + *ε* move to the harvested area with probability *α*_1_ and to the narrow band with probability 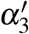. With similar methods used by Maciel and Lutscher (2013), we derived boundary conditions that both population density and flux maybe discontinuous (please see details in Appendix A). The boundary conditions can be expressed as

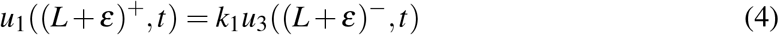

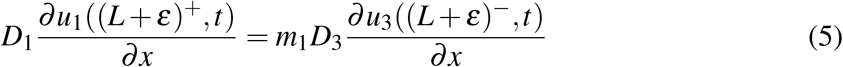

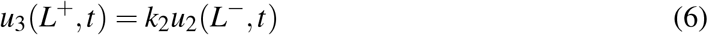

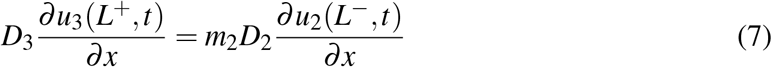

where parameters *k*_1_ and *k*_2_ represent density discontinuity at the boundaries (that is the boundary between the marine reserve and the narrow band as well as the boundary between the narrow band and the harvested area), while *m*_1_ and *m*_2_ represent flux discontinuity at the boundaries. When the coastline length of the narrow band approaches zero, the position at *L* approaches the position at *L* + *ε* and thus Eq. 4 and Eq. 6 indicate that the product (*k* = *k*_1_*k*_2_) of *k*_1_ and *k*_2_ measures the discontinuity in density between the marine reserve and the harvested area, while Eq. 5 and Eq. 7 indicate the product (*m* = *m*_1_*m*_2_) of *m*_1_ and *m*_2_ measures the discontinuous flux there. Both *k* and *m* depend on habitat preference as well as how individuals move within each patch. For example, the value of *k* will increase if individuals at the boundary move into the harvested area with high movement probability or move into the marine reserve with low movement probability; the value of *m* will increase if individuals at the boundary move into either the harvested area or the marine reserve with low movement probability (see more details in the Eq. A.21 and Eq. A.22 in Appendix A).

We assume that the environment outside the exterior boundaries (i.e. at position 0 and 1) is so hostile that individuals at the exterior boundaries cannot survive if they leave the marine reserve or the harvested area. Thus, we obtain the Dirichlet boundary conditions

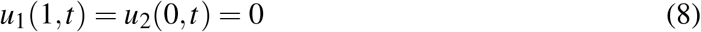

Since some individuals may lose at the boundaries, the average growth rate *λ* is smaller than the growth rate (*r*_2_) in the marine reserve where individuals grow in favorable environment. However, it is difficult to judge the relation between the average growth rate *λ* and the net growth rate (*r*_1_ − *µ*_1_*u*_1_ − *qE*_1_) in the harvested area, which should be discussed. Analytically solving the linearized form of the objective system equations under our boundary conditions, we can achieve equations that links the average growth rate *λ* and the other parameters when *ε* approaches zero (i.e. the narrow band turns into a boundary):

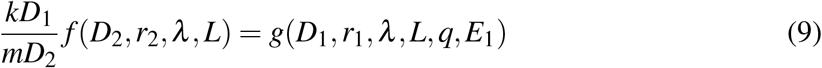

where

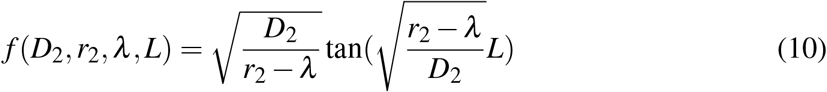

and

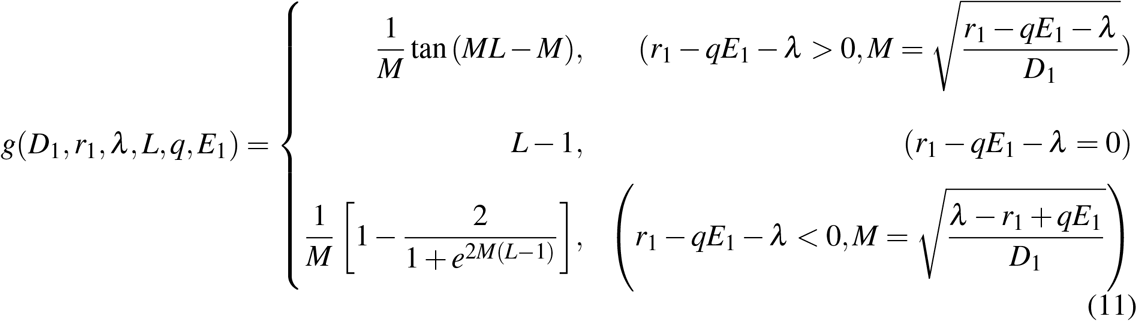

A standard calculation (see in Appendix A) indicates that *f* (*D*_2_, *r*_2_, *λ, L*) is a continuous and decreasing function of *λ*, while *g*(*D*_1_, *r*_1_, *λ, L, q, E*_1_) is a continuous and increasing function of *λ*. This property suggests that any parameter that can increase function *f* (*D*_2_, *r*_2_, *λ, L*) will increase average population growth rate *λ* when function *g*(*D*_1_, *r*_1_, *λ, L, q, E*_1_) is fixed. On the contrary, any parameter that can increase function *g*(*D*_1_, *r*_1_, *λ, L, q, E*_1_) will decrease average population growth rate *λ* when function *f* (*D*_2_, *r*_2_, *λ, L*) is fixed. Note that *k* = *k*_1_*k*_2_ and *m* = *m*_1_*m*_2_.

### Yield comparison

We further compare the harvested yields produced from the narrow band (which will be the reserve boundary when the coastline length of the narrow band *ε* approaches zero) and the harvested area. To do so, we first calculate the total yield in each region (the narrow band and the harvested area) by integrating the yield over all locations. Thus, the total harvested yields *Y*_1_ in the harvested area can be expressed as:

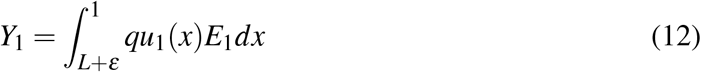

Similarly, the total harvested yields *Y*_3_ in the narrow band is:

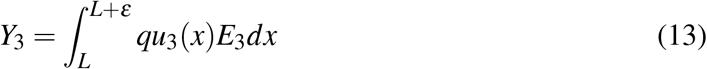

By analytically solving the population density *u*_1_ and *u*_3_, the harvested yields *Y*_1_ and *Y*_3_ can be expressed in the form with model parameters (such as population growth rate and the coastline length of the marine reserve) as well as some new mathematical constants. To cancel these mathematical constants so that we can analyze the effects of model parameters on harvested yields, we further calculate the quotient of the two kinds of yields when the coastline length of the narrow band approaches zero (see detailed derivation in Appendix A):

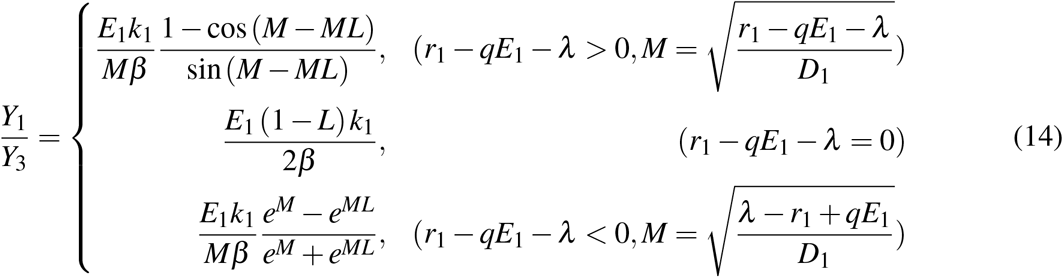

Here,through the limit outcome in Eq. 14, we can see the relative harvested yields in the harvested area in comparison to the yields at the boundary is affected by the discontinuity in density but not the discontinuity in flux although the discontinuity in flux may have effects on both types of yields separately. Generally, we would analyze how parameters regulate the value of *Y*_1_*/Y*_3_ so that to investigate the limitations and advantages of fishing the line relative to fishing in the harvested area.

### Numerical simulation

To perform simulation, we need appropriate domain for both function *f* (*D*_2_, *r*_2_, *λ, L*) and *g*(*D*_1_, *r*_1_, *λ, L, q, E*_1_). The domain used for numerical simulation corresponds to that used for analytical derivation in appendix. The domain is set to entail that the mathematical calculation is meaningful and the quotient of yields (*Y*_1_*/Y*_2_) is positive. Thus, we have 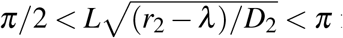 or all the three scenarios (i.e. the net growth rate in the harvested area is larger, identical to or smaller than the average population growth rate) and thus function *f* (*D*_2_, *r*_2_, *λ, L*) is negative, which leads to another condition of *r*_1_ − *qE*_1_ − *D*_1_(*π/*(2(1 − *L*)))^2^ < *λ* < *r*_1_ − *qE*_1_ for the scenario that the net growth rate in the harvested area is larger than the average population growth rate so that function *g*(*D*_1_, *r*_1_, *λ, L, q, E*_1_) is negative and Eq. 9 can hold. According to the foraging theory, we assume that individuals move slowly with small step size when they stay in the marine reserve, while individuals move fast with big step size when they stay in the harvested area, which suggests that the diffusion rate in the harvested area is bigger than that in the marine reserve (i.e. *D*_1_ > *D*_2_). Based on these conditions, we do simulations to investigate i) how growth rate in and habitat preference to the marine reserve regulate relative yields indirectly through average population growth rate? ii) how relative yields are regulated directly by average population growth rate as well as other model parameters?

## Results

The analytical derivation indicates that *f* (*D*_2_, *r*_2_, *λ, L*) is an increasing function of the growth rate (*r*_2_) in marine reserve (see details in Appendix A). Meanwhile, the parameter *r*_2_ does not exist in the function *g*(*D*_1_, *r*_1_, *λ, L, q, E*_1_), and thus variations in the growth rate in marine reserve have no effect on the function *g*(*D*_1_, *r*_1_, *λ, L, q, E*_1_). Recall that the property in Eq. 9 predicts that increasing in function *f* (*D*_2_, *r*_2_, *λ, L*) but fixing in function *g*(*D*_1_, *r*_1_, *λ, L, q, E*_1_) will increase average population growth rate *λ*. Therefore, increasing in the growth rate *r*_2_ will increase average population growth rate *λ*, and this relationship holds under three situations (i.e. the net growth rate in the harvested area is larger, identical to or smaller than the average population growth rate). On the contrary, the average population growth rate will increase if the quotient of the discontinuity in density and in flux (i.e. the term *k/m* in Eq. 9) decrease. This derives from the fact that the term *k/m* is positive while function *f* (*D*_2_, *r*_2_, *λ, L*) is negative. According to Eq. A.21 and Eq. A.22 in the Appendix A, the term *k/m* is an increasing function of the individual movement preference (*α*_2_) to the marine reserve. Therefore, high movement preference to the marine reserve will finally lead to low average population growth rate. This case can happen when fishing activities are strong and many individuals are lost at the boundary, whose negative effects on average population growth rate overwhelm the benefits from increasing movement preference to the marine reserve. The analytical results that high average population growth rate needs high growth rate in the marine reserve and low movement preference to the marine reserve are also illustrated by the simulation results (Fig. 2).

**Figure 2:**
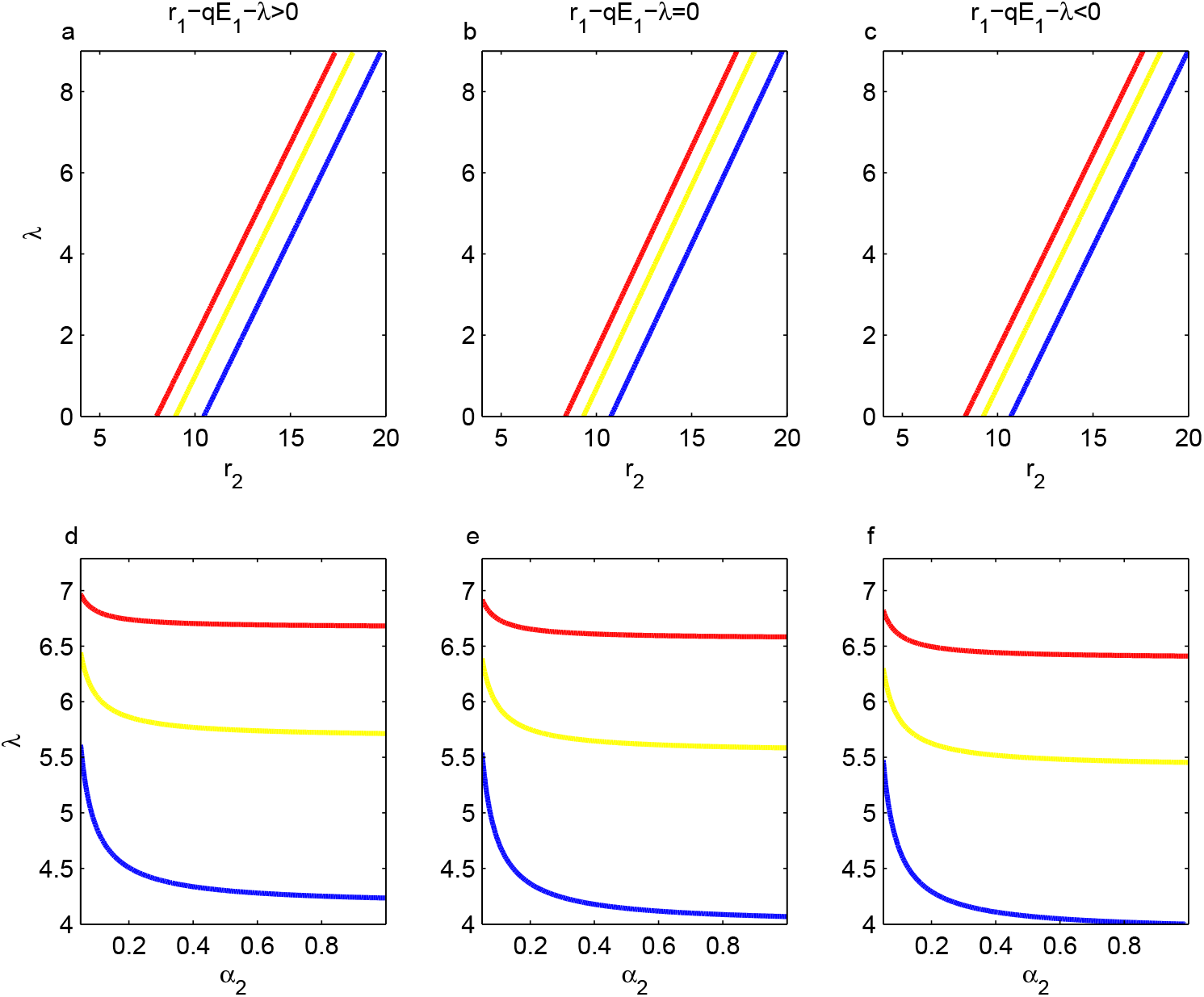
Variation of average population growth rate in response to (a-c) growth rate in marine reserve and (d-f) movement preference to reserve. Three scenarios are considered based on whether the net growth rate in the harvested area is larger, identical to or smaller than the average population growth rate. Parameter values are *D*_2_ = 0.5, *L* = 0.4, *q* = 0.1, *E*_1_ = 10, *p*_1_ = 0.6, *p*_2_ = 0.1, Δ*x*_1_ = 3, Δ*x*_2_ = 1, Δ*x*_*L*_ = 2, and *D*_1_ = 2 (red), *D*_1_ = 5 (yellow), *D*_1_ = 10 (blue). *r*_1_ = 10 (*r*_1_ − *qE*_1_− *λ* > 0), *r*_1_ = 3 (*r*_1_ − *qE*_1_ − *λ* < 0). In addition, *α*_2_ = 0.3 in (a-c) and *r*_2_ = 15 in (d-f).

Based on Eq. 14, we can easily achieve that increasing the density discontinuity (i.e. *k*_1_) at the boundary between the narrow band and the harvested area leads to high value of *Y*_1_*/Y*_3_ and thus will increase the yields benefits from fishing in the harvested area relative to fishing the line. Conversely, increasing the total fishing efforts (i.e. *β*) in the narrow band will increase the relative benefits of yields from fishing the line. The two important relationships hold under three situations (i.e. the net growth rate in the harvested area is larger, identical to or smaller than the average population growth rate). Moreover, further analytical (see details in Appendix A) and simulations results indicate that increasing the average population growth rate as well as the coastline length of the marine reserve but decreasing the growth rate outside the marine reserve will increase the relative benefits of yields from fishing the line (Fig. 3), and the relationships hold regardless of whether the net growth rate in the harvested area is larger, identical to or smaller than the average population growth rate. Note that the average population growth rate can indirectly increase the relative benefits of yields from fishing the line although Eq. 14 shows that average population growth rate does not regulate the relative yields (i.e. the quotient of the yields from harvested area and the yields from fishing the line) when the net growth rate in the harvested area is identical to the average population growth rate. This derives from the fact that, on such a case, the fishing effort per unit length in the harvested area has a positive relationship with the quotient of the yields from harvested area and the yields from fishing the line but a negative relationship with the average population growth rate.

**Figure 3:**
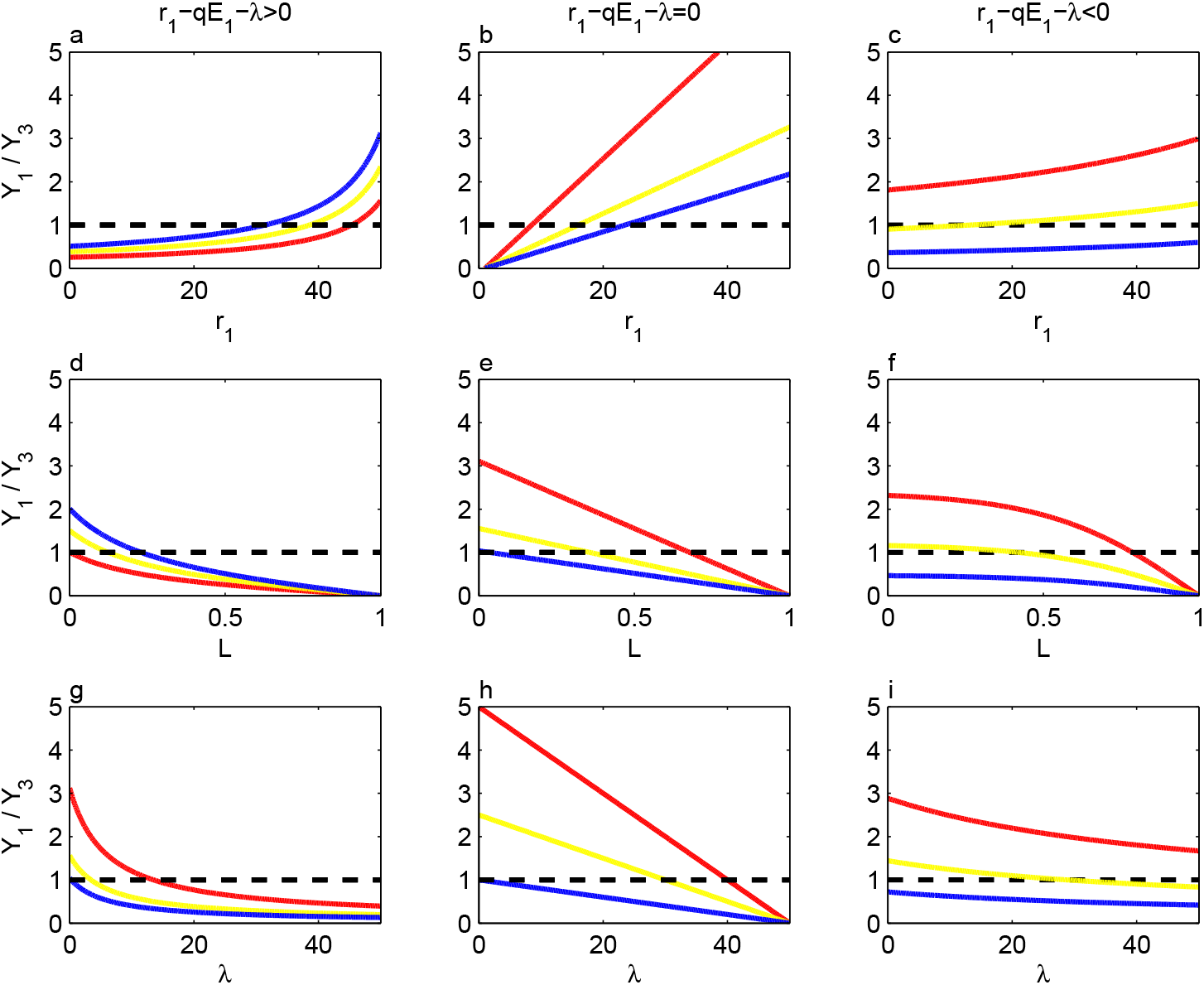
Variation of relative yields in response to (a-c) growth rate outside the reserve, (d-f) the coastline length of marine reserve and (g-i) average population growth rate. Parameter values for all subplots are *D*_1_ = 2, *q* = 0.1, *p*_1_ = 0.6, *p*_2_ = 0.1, *p*_3_ = 0.3, *α*_1_ = 0.2, *α*_2_ = 0.3, Δ*x*_1_ = 3, Δ*x*_2_ = 1, Δ*x*_3_ = 2, Parameter values for each subplot are (a,d) *β* = 1.5; *E*_1_ = 10 and *λ* = 2 (red), *E*_1_ = 15 and *λ* = 1.5 (yellow), *E*_1_ = 20 and *λ* = 1 (blue), (b,e) *λ* = 1; *β* = 3 (red), 6 (yellow) and 9 (blue), (c,f) *λ* = 20, *E*_1_ = 300; *β* = 4 (red), 8 (yellow) and 20 (blue), (g) *E*_1_ = 10, *r*_1_ = 50; *β* = 1 (red), 2 (yellow) and 3 (blue), (h) *β* = 4 (red), 8 (yellow) and 20 (blue), (i) *E*_1_ = 100; *β* = 1.5 (red), 3 (yellow) and 6 (blue). In addition, *L* = 0.4 in (a-c, g-i) and *r*_1_ = 15 in (d-f, i).

Our results have shown that decreasing the density discontinuity *k*_1_ and increasing the total fishing efforts *β* can increase the relative benefits of yields from fishing the line. This has important implications in fisheries management as the density discontinuity *k*_1_ is an decreasing function of the movement preference 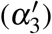 to the boundary between the marine reserve and the harvested area. It suggests that increasing the movement preference to the boundary between the marine reserve and the harvested area will finally lead to increasing in the relative benefits of yields from fishing the line. A strong fishing activities in combination with fish individual’s high movement preference to the reserve boundary will finally lead to large amount of fish individuals losing at the boundary between the marine reserve and the harvested area. Therefore, the advantages of fishing the line in harvesting yields depends on fish individual losing at the reserve boundary. When the coastline length of the narrow band *ε* approaches zero, the sum of the probability of the movement preference to the marine reserve (*α*_2_), at the boundary 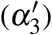 and to the harvested area (*α*_1_) is 1, which suggests that density discontinuity *k*_1_ and thus the relative yields (*Y*_1_*/Y*_3_) increase with high movement preference to the marine reserve (*α*_2_). Although the relative yields does not directly regulated by the growth rate in the marine reserve according to Eq. 14, it may be influenced indirectly through changing average population growth rate. Our results have shown that high growth rate in the marine reserve will increase average population growth rate (see details in Appendix A) which will lead to increasing in the relative benefits of yields from fishing the line. On the contrary, the direct analytical results from Eq. 14 indicate that growth rate outside the reserve leads to reverse effects (see appendix). Thus, the advantages of fishing the line in harvesting yields depends on a high growth rate inside and low growth rate outside the marine reserve as well. An important goal in ecological conservation is maintaining population persistence rather than over-fishing. To achieve this goal, the average population growth rate should not be too low, which can ensure a high growth rate in the marine reserve and thus the advantages of fishing the line emerge.

## Discussion

With suitable boundary conditions, we have found that fishing the line should not be the permanent strategy if fishermen want to improve their harvested yields. The advantages of fishing the line in harvested yields can only emerge if marine reserves can provide great benefits so that the target fish population has a high growth rate inside a marine reserve relative to the growth rate outside the reserve. The advantages will be strengthened by individual high movement preference to the boundary and by increasing marine reserve size. Our findings have very important implications of not using harvesting tactic of fishing the line in terms of new established marine reserves in which population might have a very low growth rate because of over-fishing in history. Meanwhile, our findings suggest that fishing the line in terms of a small marine reserve is the worst strategy as neither the goal of maximizing harvested yields nor the goal of conservation in the target fish species can be achieved. Kellner et al. (2007) studied the optimal effort distribution in the harvested area and at habitat boundary, which may economically inspire fishers to spend less cost and achieve more yields benefits. Their findings suggest that the optimal strategies in harvested yields must contain the components of fishing the line. However, our results have shown that fishing the line cannot exhibit the advantages in harvested yields under some conditions (such as individuals have a low growth rate in a small marine reserve) even if the fishing effort per unit length at the reserve boundary is extremely strong (approach infinite) which should not be an economic way in fisheries. The different conclusions between ours and previous ones may be derived from the fact that our theoretical framework includes more reasonable and realistic features by considering individual movement behavior and individual losing at habitat boundaries. Without habitat preference, Kellner et al. (2007) have shown that population density in the marine reserve decrease if individual movement rate increase, which may not happen if individuals have high movement preference to the marine reserve simultaneously [19]. Moreover, considering habitat preference and discontinuity at the reserve boundary, effects of individual movement rates are different between the marine reserve and the harvested area. Indeed, our results indicate that increasing movement rate in the harvested area or decreasing movement rate at the reserve boundary can enhance relative benefits of yields from fishing the line (see detail in Appendix A). Note that the intuitive reasonable effect of movement rate may not be general as we can only draw conclusions under special cases by assuming that either movement probabilities or step size are equal between the harvested area and the reserve boundary.

### Assumptions

One of the assumptions we used is that fish individuals are self-replenishment in the harvested area, which suggests that the net growth rate (i.e. *r*_1_ − *qE*_1_) in the harvested area is positive without the necessity of spillover from the marine reserve to maintain population persistence. On such a case, the uncertain relationship between the net growth rate in the harvested area and the average population growth rate increase the complexity of the scientific problem we focus on here. For example, increasing fishing effort per unit length in the harvested area can increase or decrease the relative yields in some cases while has a monotonous effect on the relative yields in other cases (see detail in Appendix A). If the net growth rate in the harvested area is negative, analyzing the advantages of fishing the line become simple but adding population persistence conditions (the coastline length of the marine reserve should not be smaller than a critical value to maintain persistence) because of the source-sink relationship between the marine reserve and the harvested area. The negative net growth rate in the harvested area will always smaller than the average population growth rate (should be positive to maintain population persistence), which leads to a clear increasing effect of fishing effort per unit length in the harvested area on the quotients of yields from the harvested area and from fishing the line. The persistence condition of minimum marine reserve size suggests that, for some special cases without changing other parameters, variation of the marine reserve size can always lead to higher harvested yields from fishing the line than that from the harvested area because our analytical results have shown that increasing marine reserve size increases the relative benefits of yields from fishing the line.

Another assumption in the paper is based on the foraging theory, which suggests that individuals have high probability moving into favorable patches but move slowly once they arrive there. Thus, we assume that individuals move fast to the marine reserve with big step size but move slowly when they stay in the marine reserve, while individuals move slowly toward the harvested area with small step size but move fast when they stay in the harvested area. Therefore, the value of discontinuous flux (i.e. *m*) is positive, which leads to an increasing relationship between function *f* (*D*_2_, *r*_2_, *λ, L*) and the left side of Eq. 9. If the foraging theory does not hold and the assumptions of movement preference are changed, the value of discontinuous flux may be negative, which will further lead to the reverse in most of the results we achieved in the paper.

In addition, the dependencies and trade-offs among the parameters in Eq. 9 and Eq. 14 may change the outcome of our objective system. Under constant losing effect at the boundary (i.e. the total harvested effort *β* is constant), individuals at the boundary may increase the preference to move into the marine reserve if fishing activities is too strong in the harvested area. In other words, individuals movement probability (*α*_2_) to the marine reserve depends on the difference (*r*_2_ − (*r*_1_ −*qE*_1_)) in habitat quality (here, it is denoted by growth rate because high habitat quality corresponds to high growth rate, and low habitat quality corresponds to low growth rate.) between the favorable marine reserve and the unfavorable harvested area. This relationship can be expressed by a classic equation [20], and the reanalysis results indicate that the conclusion (i.e. large marine reserve size with high individual growth rate inside and low growth rate outside, high movement preference to the marine reserve as well as individual losing at boundary jointly promote the advantages of fishing the line) does not change on such a case.

### Further applications of boundary condition

The universal phenomenon in ecology is that habitat heterogeneous which will lead to habitat boundaries in both terrestrial and marine systems. Habitat at boundaries may be more suitable for individual to live because the environment there possess large amounts of the characteristics of different habitats nearby. Therefore, different species moves into the habitat boundaries including the predator which will cause the losing of the prey at boundaries. A classical example is the so-called “ecological trap hypothesis”, which believes that a number of birds populations prefer to reproduce at habitat boundary where mortality is relatively high (individuals losing at the boundary) because the individuals there are much easier to be found by their natural enemies [21]. Therefore, the phenomenon that losing individuals at habitat boundaries is much universal in ecosystem, which is similar to the phenomenon of fishing the line and the boundary condition we derived here can be applied to the similar situations.

An important example is applying the boundary condition with discontinuous flux into the refuge design, which achieves more reasonable conclusions than previous ones. Cantrell and Cosner (1999) have investigated a landscape with favorable refuge and unfavorable buffer zone. Their results have shown that high quality in the unfavorable buffer zone decreases the average population growth rate, which finally needs to increase the minimum length of favorable refuge to maintain population persistence. The explanation for the counter-intuitive phenomenon is that increasing quality in the buffer zone will increase individuals moving into hostile surroundings from the buffer zone such that increase individual mortality. On the contrary, individuals will stay in the favorable refuge because of the aversion to the buffer zone created by the poor quality. Using boundary conditions with discontinuous density and continuous flux, Maciel and Lutscher (2013) drew the same conclusion based on the assumption that individuals leaving one patch will enter another and cannot lose or add at habitat boundaries. However, with our boundary condition, our analysis results indicate that high quality in the buffer zone leads to high average population growth rate (see detail in Appendix B) and thus the counter-intuitive phenomenon disappears. This may be derived from the fact that considering individual losing at habitat boundaries is much more reasonable in the real-world system.

### Future directions

A widely accepted concept is that marine reserves can enhance fish individual’s habitat quality such as shelter or food availability through restricting destructive fishing activities while over-fishing leads to habitat degradation in the harvested area [6, 19]. On such a case, high growth rate inside while low growth rate outside the marine reserve will be much common. Supposing a species is so sensitive to the habitat quality that it cannot survive when the habitat quality only decreases a bit, then the trends that high growth rate inside and low outside the reserve should be strengthened. Therefore, species with different sensitivities to the habitat quality result in variation of the discrepancy between growth rate inside and outside the reserve, which suggests that the optimal solution exhibiting advantages of fishing the line for one species may not be optimal for another species. Further research should pay much attention to investigate the harvested yields advantages of fishing the line for a strong stock (which has strong viability and is less sensitive to habitat quality) under the persistence of a weak stock (which is much more sensitive to the habitat quality and easily goes extinct).

## Appendix A

### Mathematical Derivation and Analysis

#### Discontinuous boundary condition

Similar to Maciel & Lutscher (2013), we derive the discontinuous boundary conditions by taking both random walk and habitat preference into consideration. Individuals movement in the same habitat belongs to random walk (i.e. *p*_*i*_ in patch *i*), which means movement probability to both direction (left or right) are the same. Individuals movement in different habitats belongs to habitat preference 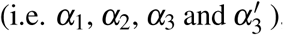, which means movement probability at the boundary are different towards different habitats. We assume that individuals move along straight line in discrete time steps. The step size is Δ*x*_*L*_ at position *L*, Δ*x*_*L*+*ε*_ at position *L* + *ε* and Δ*x*_*i*_ in patch *i* (Again, recall that *i* = 1 for harvested area, *i* = 2 for marine reserve, *i* = 3 for narrow band). We assume that the probability function at time *t* is *P*(*x, t*), where *x* means individuals at positions *L* − Δ*x*_2_, *L, L* + Δ*x*_3_, *L* + *ε* − Δ*x*_3_, *L* + *ε, L* + *ε* + Δ*x*_1_, respectively. We first derive the relationships that are applicable to the boundary between the marine reserve and the narrow band:

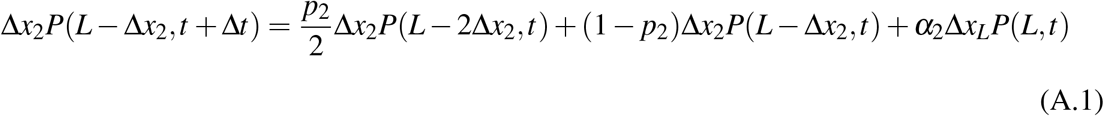

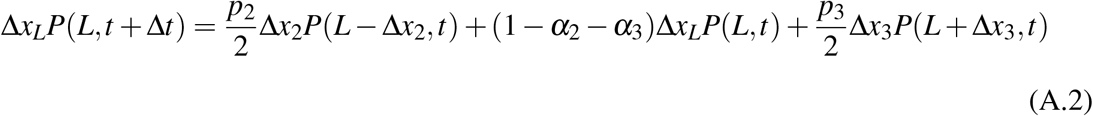

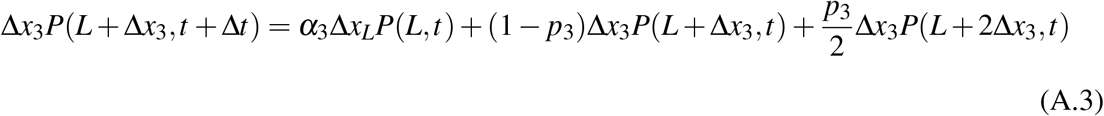

According to Taylor series, the above equations can be approximated as:

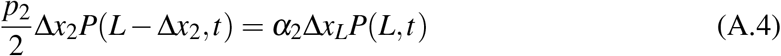

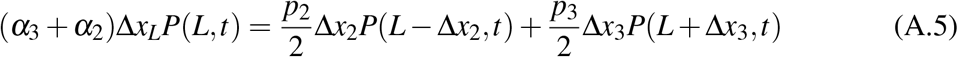

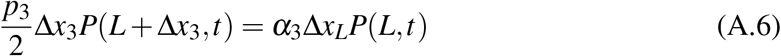

In combination with equations 4 and 6, we have:

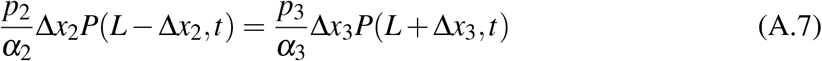

Let 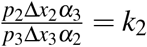, then:

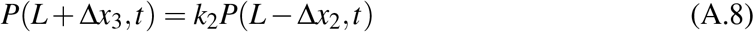

Substitute the probability density function with density:

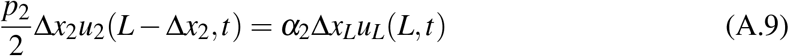

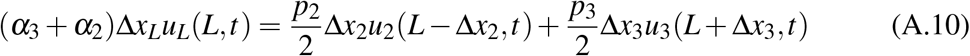

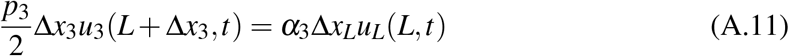

Therefore, we have:

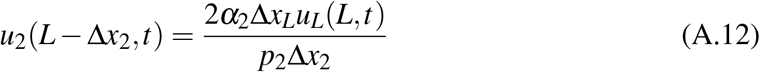

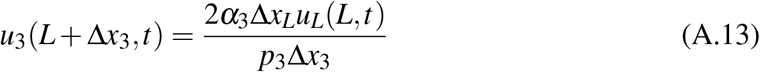

The flux has the following relationship:

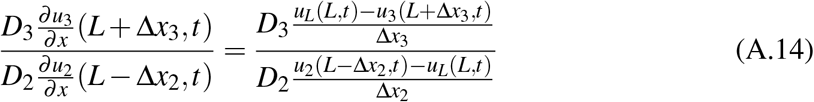

Like previous research (Maciel & Lutscher 2013), the diffusion coefficient is defined as 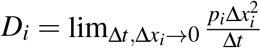. Substitute Eqs 12 and 13 into 14, we have:

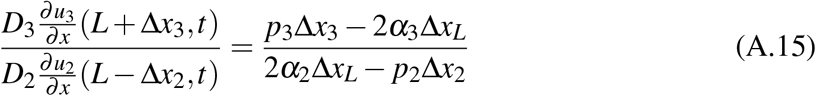

Let 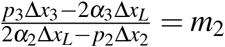, then:

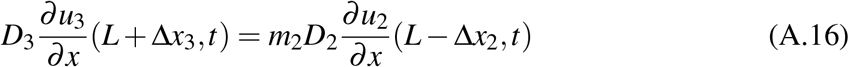

Therefore, equation 15 or 16 is the discontinuous flux boundary condition when constant *m*_2_ not equals 1. Using the same method to get the relationships for the boundary between the narrow band and the harvested area:

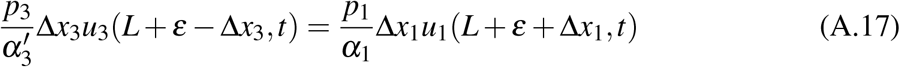

i.e.

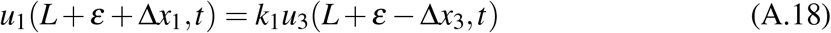

where 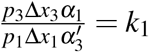. Similarly, we have the flux relationship:

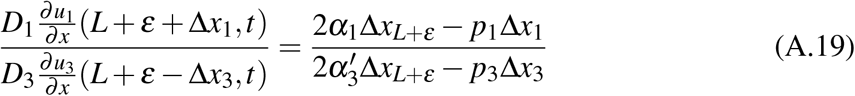

i.e.

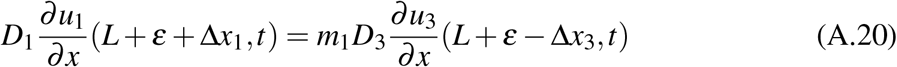

where 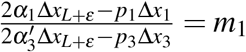. When the coastline length of the narrow band (*ε*) approaches zero, the position *x* = *L* + *ε* will be the position *x* = *L*, which suggests that individuals at both positions (*x* = *L* + *ε* and *x* = *L*) have the same probability of moving into the narrow band (i.e. Δ*x*_*L*+*ε*_ = Δ*x*_*L*_, *α*_3_ = *α*3^*t*^). Therefore, the product (*k*) of *k*_1_ and *k*_2_ as well as the product (*m*) of *m*_1_ and *m*_2_ will be

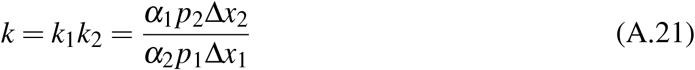

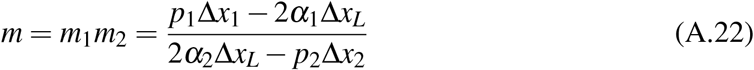

According to the foraging theory, we assume that individuals move fast with big step size in the harvested area while have low preference and small step size at the boundary toward the harvested area, and the situation is reversed for individuals in the marine reserve. Therefore, the value of *m* is positive.

#### Effect of model parameters on average population growth rate

Based on the exponential solutions *u*_*i*_(*x, t*) = *e*^*λt*^*V*_*i*_(*x*), (*i* = 1, 2, 3), the equations 1, 2 and 3 in the main text can be transformed into:

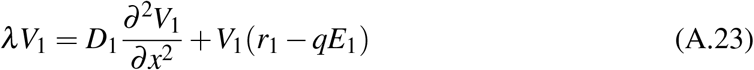

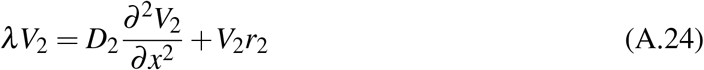

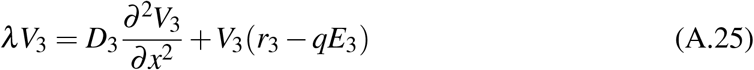

thus, the solutions of *V*_1_, *V*_2_, *V*_3_ are

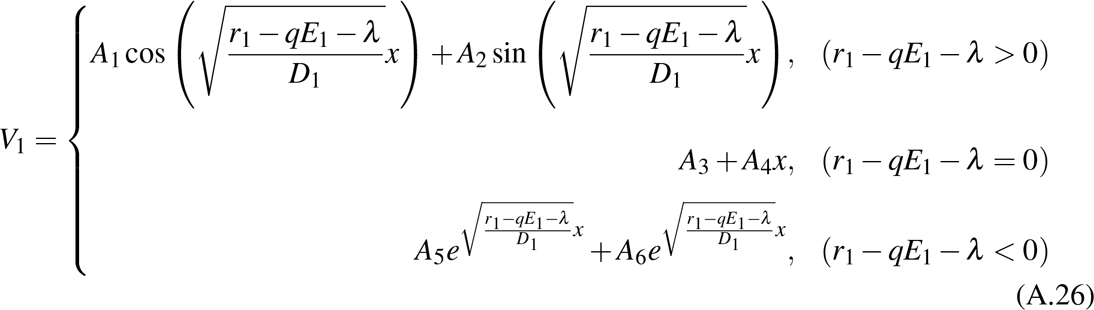

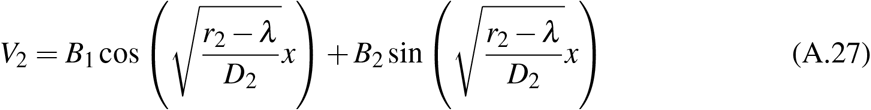

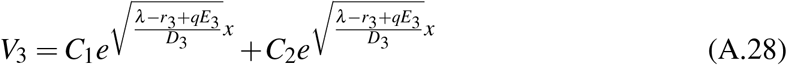

where *A*_1_ to *A*_6_, *B*_1_, *B*_2_, *C*_1_ and *C*_2_ are constants, and the boundary conditions are

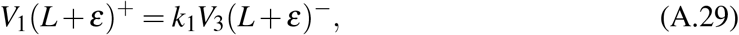

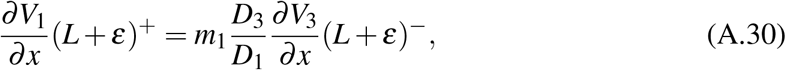

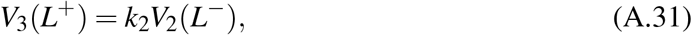

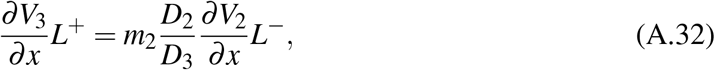

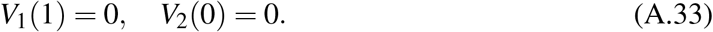

Using the solutions of *V*_1_, *V*_2_, *V*_3_, we can achieve their derivations. Substitute the solutions of *V*_1_, *V*_2_, *V*_3_ as well as their derivations into the boundary conditions, and let 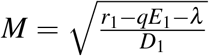 and 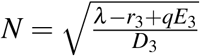, we have six relationships under the condition of *r*_1_ −*qE*_1_ −*λ* > 0

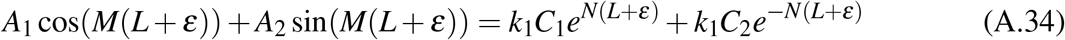

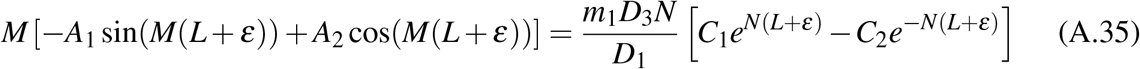

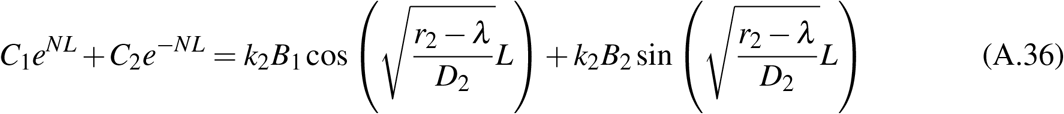

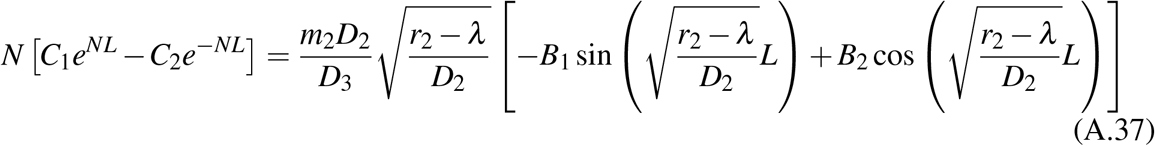

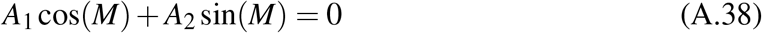

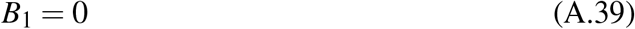

when *ε* approaches zero, this leads to

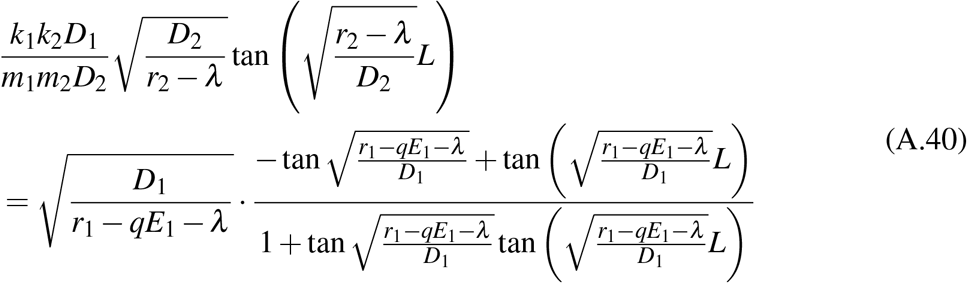

Similarly, when *r*_1_ −*qE*_1_ −*λ* = 0 and *ε* approaches zero, we have

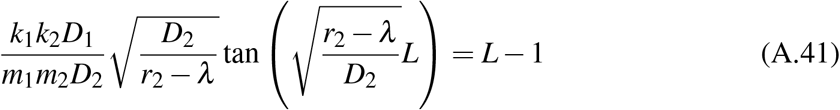

when *r*_1_ −*qE*_1_ −*λ* < 0 and *ε* approaches zero, we have

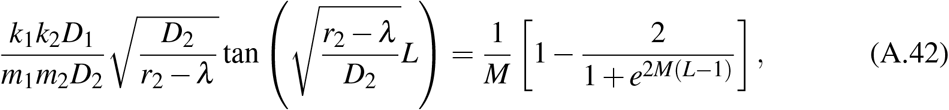

where 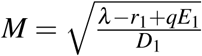 (Note that we denote 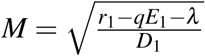 when *r*_1_ −*qE*_1_ −*λ* > 0). In summary, the relationship between model parameters and the average population growth rate (*λ*) can be expressed as

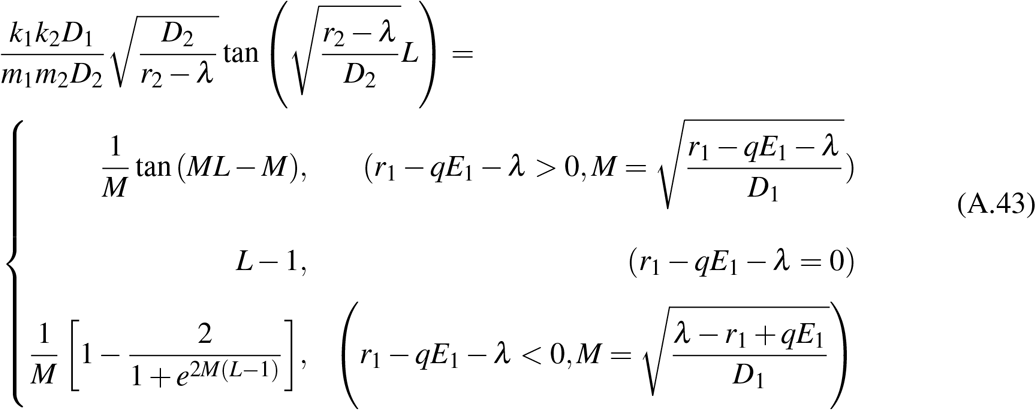

Let

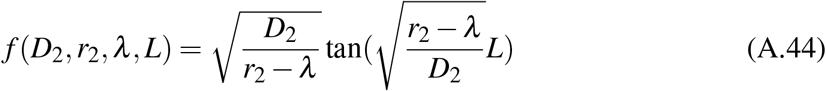

and

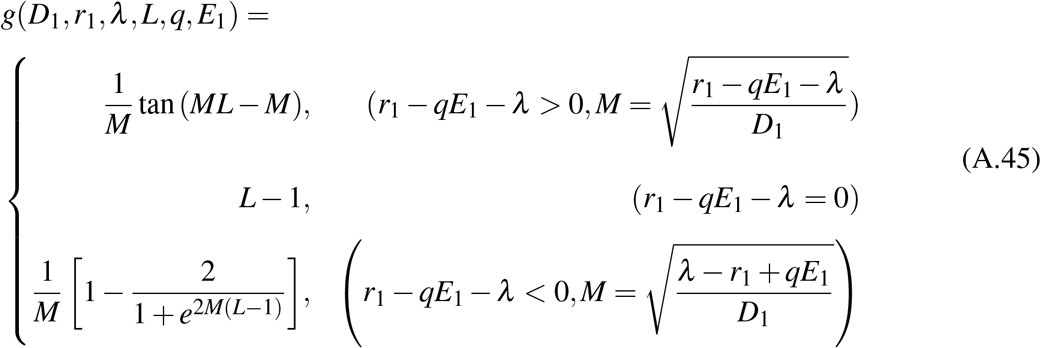

To investigate the monotonicity of function *f* (*D*_2_, *r*_2_, *λ, L*) and *g*(*D*_1_, *r*_1_, *λ, L, q, E*_1_), we have the derivation

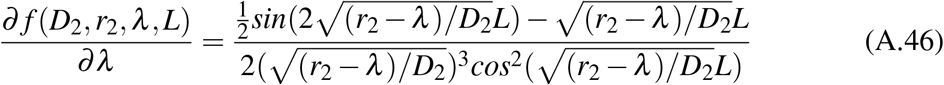

let 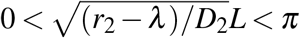, then 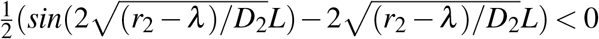. Thus, *∂ f* (*D*_2_, *r*_2_, *λ, L*)*/∂λ* < 0, and function *f* (*D*_2_, *r*_2_, *λ, L*) is decreasing in *λ*. When *r*_1_ −*qE*_1_ −*λ* > 0, we have

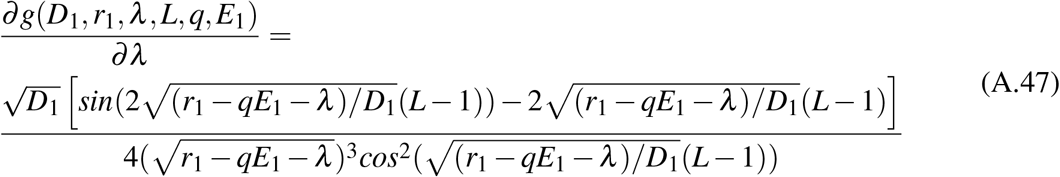

Because *L* − 1 < 0, we have 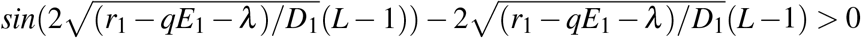, and thus *∂g*(*D*_1_, *r*_1_, *λ, L, q, E*_1_)*/∂λ* > 0. Similarly, when *r*_1_ −*qE*_1_ −*λ* < 0, we have

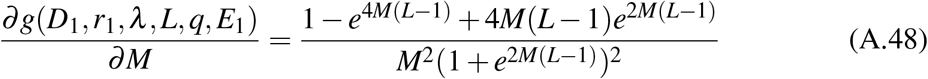

where 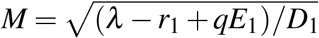. Let *F*(*M*) = 1 −*e*^4*M*(*L*−1)^ + 4*M*(*L*− 1)*e*^2*M*(*L*−1)^, then the derivation *F*^*t*^(*M*) > 0 and thus *F*(*M*) > *F*(0) = 0 and *∂g*(*D*_1_, *r*_1_, *λ, L, q, E*_1_)*/∂M* > 0. Moreover, *M* is increasing in *λ*. In summary, function *g*(*D*_1_, *r*_1_, *λ, L, q, E*_1_) is increasing in *λ* for both scenarios (i.e. *r*_1_ − *qE*_1_ − *λ* > 0 or *r*_1_ − *qE*_1_ − *λ* < 0). We further demonstrated that function *f* (*D*_2_, *r*_2_, *λ, L*) is increasing in *r*_2_ by calculating the derivation

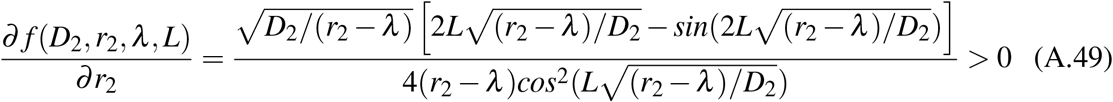

#### Yield comparison

According to the solution of *V*_3_, we can calculate the integral of *V*_3_ and thus the total harvested yield at the boundary (*Y*_3_) can be expressed as

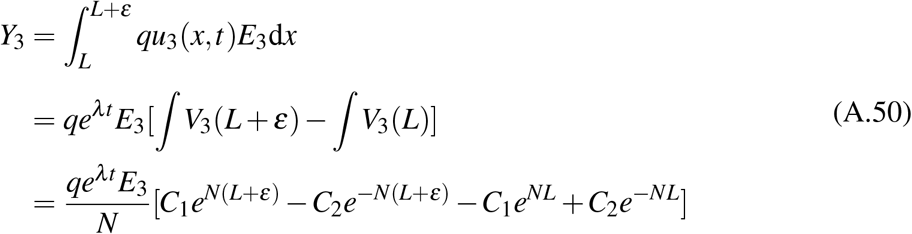

Similarly, when *r*_1_ −*qE*_1_ −*λ* > 0, the total harvested yield in the harvested area (*Y*_1_) are

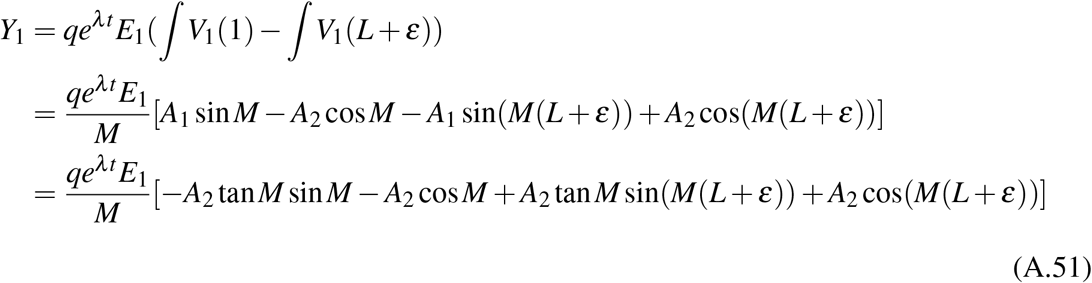

According to the boundary conditions, we have

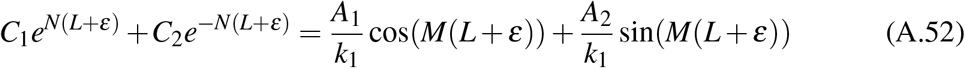

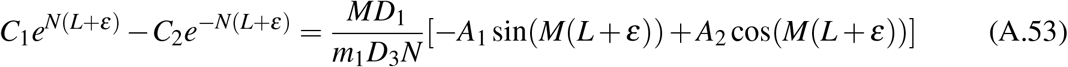

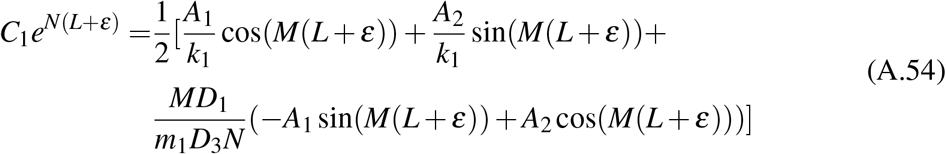

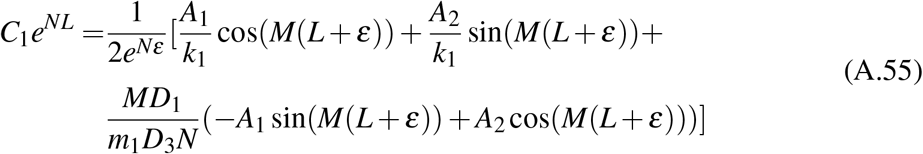

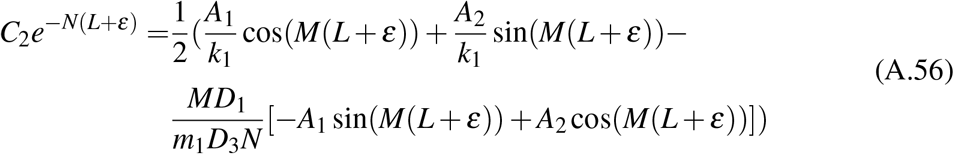

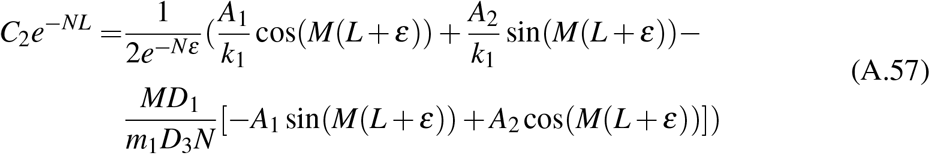

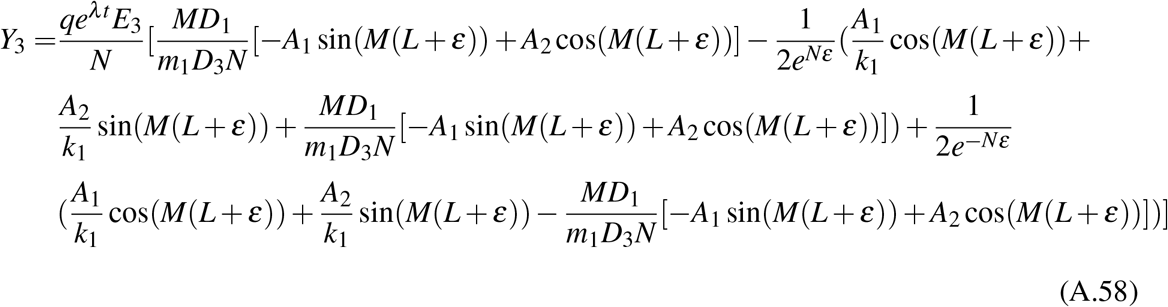

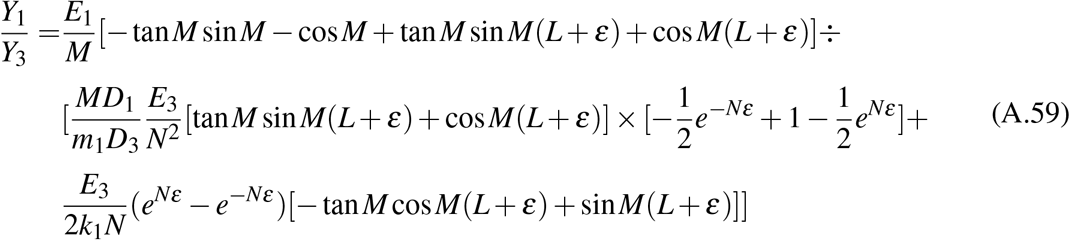

Using Taylor’s formula to calculate the limit when *ε* approaches zero, then we have

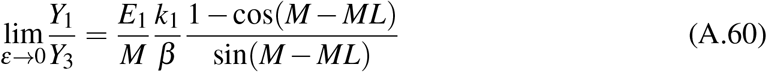

Similarly, when *r*_1_ −*qE*_1_ −*λ* = 0, the quotient of *Y*_1_ and *Y*_3_ is

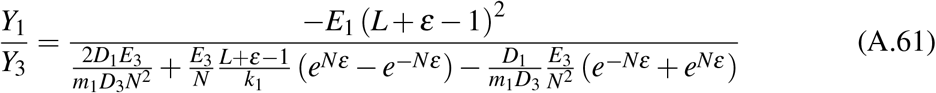

When *ε* approaches zero, the limit of *Y*_1_*/Y*_3_ is

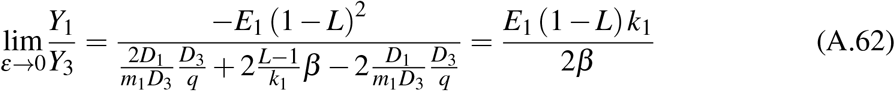

when *r*_1_ −*qE*_1_ −*λ* < 0 and *ε* approaches zero, the limit of the quotient of *Y*_1_ and *Y*_3_ is

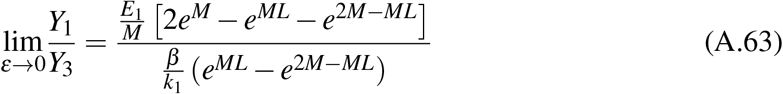

In summary, when *ε* approaches zero, the limits of the quotient of *Y*_1_ and *Y*_3_ are

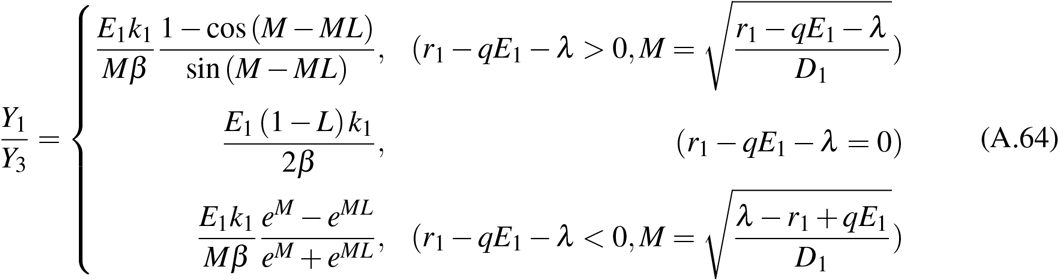

We discuss how *Y*_1_*/Y*_3_ varies with *r*_1_ and *λ*. When *r*_1_ −*qE*_1_ −*λ* > 0, we have

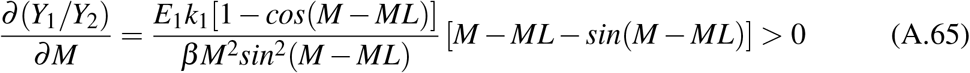

Because *M* is increasing in *r*_1_ and decreasing in *λ, Y*_1_*/Y*_3_ will increase in *r*_1_ but decrease in *λ*. When *r*_1_ −*qE*_1_ −*λ* < 0, we have

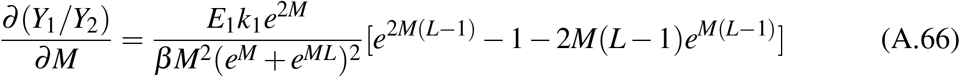

Let *θ* = *M*(*L* − 1) and *F*(*θ*) = *e*^2*θ*^ − 1 − 2*θe*^*θ*^, then *F*^*′′*^(*θ*) < 0. The negative value of *θ* makes that *F*^*′*^(*θ*) > *F*^*′*^(0) = 0. Thus, *F*(*θ*) < *F*(0) = 0 and *∂* (*Y*_1_*/Y*_2_)*/∂M* < 0, which suggests that *Y*_1_*/Y*_3_ is decreasing in *M*. Because *M* is decreasing in *r*_1_ and increasing in *λ* when *r*_1_ − *qE*_1_ −*λ* < 0, *Y*_1_*/Y*_3_ will increase in *r*_1_ but decrease in *λ*. In summary, *Y*_1_*/Y*_3_ increases in *r*_1_ but decreases in *λ* for both two scenarios (i.e. *r*_1_ − *qE*_1_ − *λ* > 0 or *r*_1_ − *qE*_1_ − *λ* < 0). Similarly, the decreasing relationship between *Y*_1_*/Y*_3_ and *L* is demonstrated by the derivation

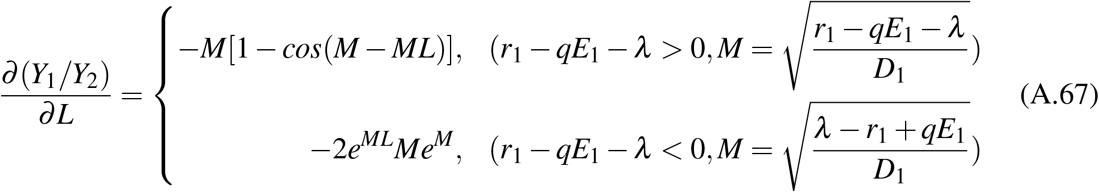

We discuss the effect of diffusion rate (*D*_1_) in the harvested area on relative yields (*Y*_1_*/Y*_3_) under two special cases: i) individual step size in the harvested area is the same to that at the boundary (i.e. Δ*x*_3_ = Δ*x*_1_); ii) individual movement rate in the harvested area is the same to that at the boundary (i.e. *p*_3_ = *p*_1_). For the first case, 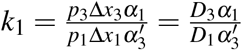, and thus we have

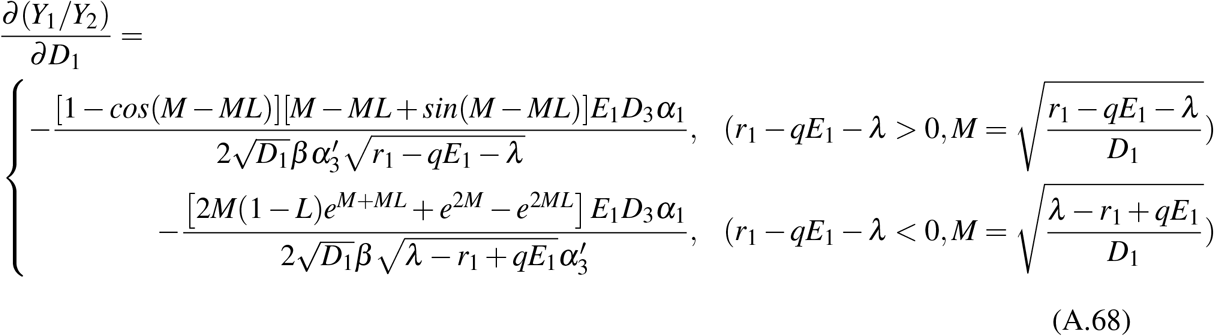

which turns out to be negative. For the second case,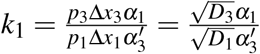, and thus we have

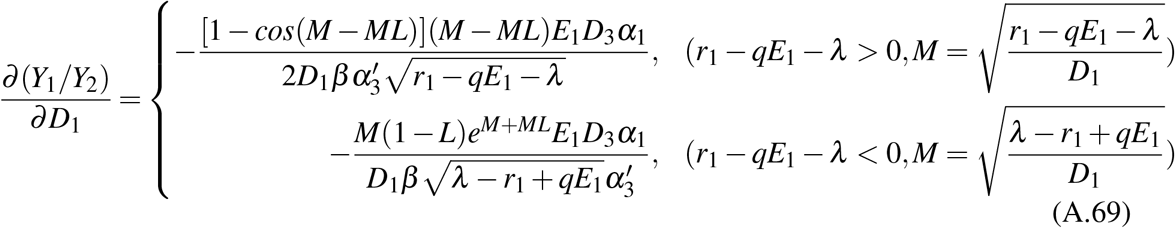

which is also negative. We can, therefore, summarize that the relative yields (*Y*_1_*/Y*_3_) increase with the decreasing of diffusion rate in the harvested area (*D*_1_) under special cases.

## Appendix B

### Applications to refuge design

Following Cantrell and Cosner (1999) as well as Maciel and Lutscher (2013), we focus on the system that consists of the favorable core habitat and the unfavorable buffer zone, and thus population growth rate in the core habitat (*r*_1_) is bigger than that in the buffer zone (*r*_2_). The system is simplified as one spatial dimension with favorable patch (i.e. core habitat) size *L*_1_ and unfavorable patch (i.e. buffer zone) size *L*_2_. Supposing position zero is the boundary between favorable and unfavorable patch, we have the model (Cantrell and Cosner 1999; Maciel and Lutscher 2013)

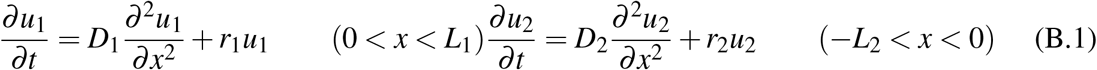

where *u*_*i*_ and *D*_*i*_ are density and diffusion rate in patch *i* (*i* = 1 for core habitat and *i* = 2 for buffer zone). With the solutions of the form *u*_*i*_(*x, t*) = *e*^*λt*^*V*_*i*_(*x*), we achieve

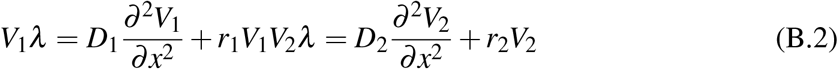

The exterior boundaries at position −*L*_2_ and *L*_1_ are the same to that in the work of Cantrell and Cosner (1999) as well as Maciel and Lutscher (2013), and thus we have *u*_2_(−*L*_2_, *t*) = 0 and (*∂u*_1_*/∂x*)(*L*_1_, *t*) = 0. However, the boundary conditions between core and buffer are different and can be shown as

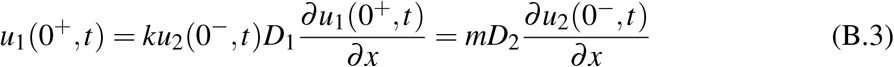

To compare the effect of our boundary conditions with previous ones, we firstly solve the analytic solutions of *V*_1_ and *V*_2_. The solutions of *V*_1_ and *V*_2_ are

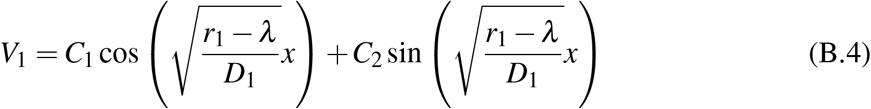

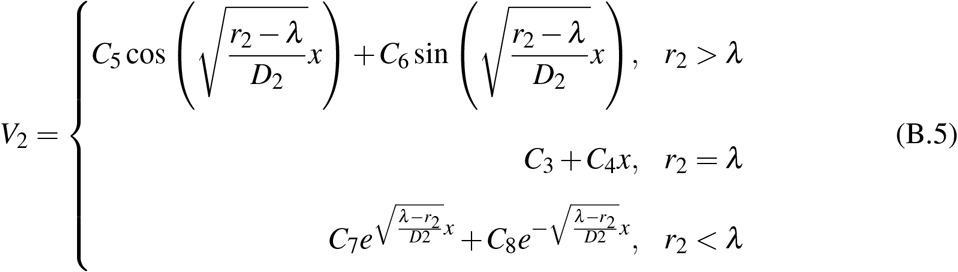

Based on the solutions of *V*_1_ and *V*_2_, we can calculate their derivations. Substitute the solutions and derivations of *V*_1_ and *V*_2_ into the boundary condition to achieve the relationships between model parameters and average growth rate. When *r*_2>_*λ*, we have the relationships

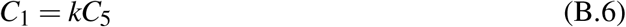

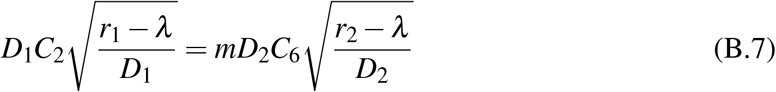

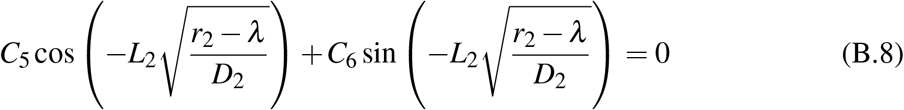

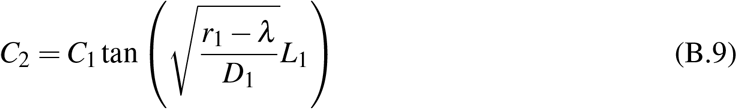

Removing constants *C*_1_, *C*_2_, *C*_5_, *C*_6_ to achieve the relationship

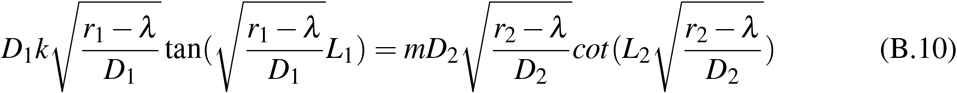

Similarly, when *r*_2_ = *λ*, the relationship is

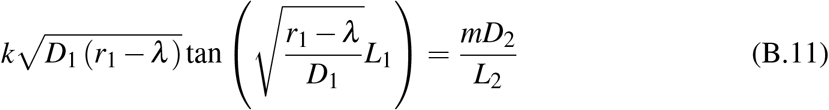

when *r*_2_ < *λ*, the relationship is

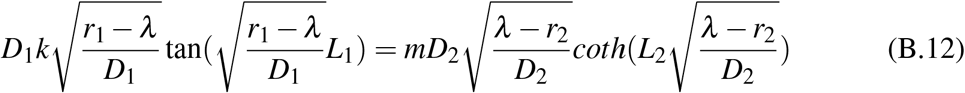

The three relationships can turn into

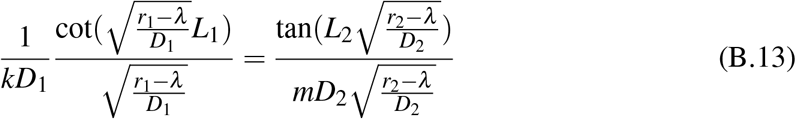

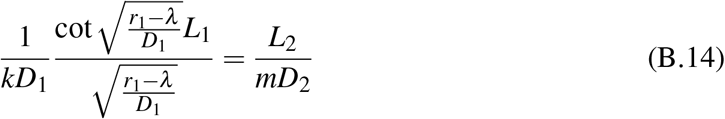

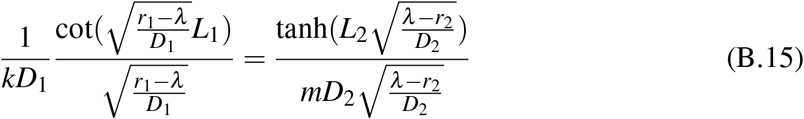

Therefore, the relationship between model parameters and average growth rate can be summarized as

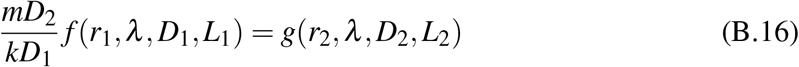

where

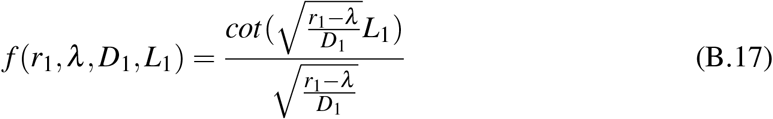

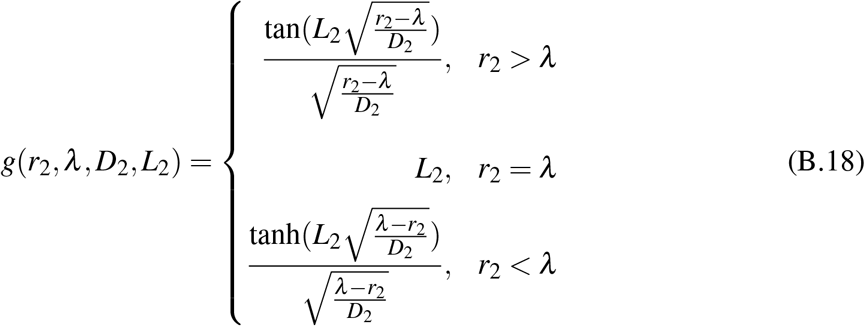

According to Cantrell and Cosner (1999), the function *f* (*r*_1_, *λ, D*_1_, *L*_1_) and thus the function 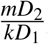 *f* (*r*_1_, *λ, D*_1_, *L*_1_) (because there is no parameter *λ* in the positive term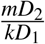) is increasing in *λ* while function *g*(*r*_2_, *λ, D*_2_, *L*_2_) is decreasing in *λ* (which actually can be achieved by calculating the derivation of function *f* (*r*_1_, *λ, D*_1_, *L*_1_) and *g*(*r*_2_, *λ, D*_2_, *L*_2_) in *λ*). Based on the calculation of the discontinuous flux boundary condition (i.e. 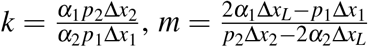. Note that *m* is positive based on the foraging theory), the term 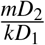 is an increasing function of *α*_1_ (i.e. movement preference to the core habitat). Therefore, the increasing of movement preference to the core habitat corresponds to the decreasing of average growth rate *λ*. This case may happen when large individuals are lost at the boundary, whose negative effect on average growth rate overwhelm the benefit from increasing preference to the core habitat. Further, by considering the dependence between habitat preference and habitat quality, we can achieve that the increasing of habitat quality in the buffer zone will lead to the increasing of average growth rate.

